# A massively parallel reporter assay of *MECP2* cis-regulatory elements reveals genetic candidates for male-biased autism

**DOI:** 10.64898/2026.05.08.723809

**Authors:** Rebecca Meyer-Schuman, Fisher Cherry, Yang Sui, Athanasios Papastathopoulos-Katsaros, Yi Zhong, Yidan Li, Tianyun Wang, Kelsey Hennick, Druha Karunakaran, Hanna Berk-Rauch, Zhandong Liu, Aravinda Chakravarti, Tomasz Nowakowski, Evan E. Eichler, Huda Zoghbi

## Abstract

Autism affects males four times more often than females, yet the basis of this sex bias remains unclear. One hypothesis is that hypomorphic variants in X-linked genes—genes where loss-of-function alleles cause syndromic neurodevelopmental disorders (NDDs) predominantly in females—produce milder, non-syndromic phenotypes in hemizygous males. We tested this by investigating cis-regulatory elements (CREs) of *MECP2*, a dosage-sensitive X-linked gene. Using a massively parallel reporter assay in human neurons, we mapped transcription factor binding sites within *MECP2* CREs and tested autism-associated variants for functional impact. We identified two noncoding variants that change CRE activity, each with a male-biased phenotype. One of these, a promoter variant, disrupts NFY binding and reduces *MECP2* expression by ∼30%, a magnitude that produces autism-like phenotypes in mice. These findings suggest noncoding *MECP2* variants can cause non-syndromic, male-biased autism, and provide a framework for uncovering regulatory variants in other X-linked NDD genes that may contribute to autism’s missing heritability.

## Introduction

Autism is a common, heritable^1^ neurodevelopmental disorder (NDD) marked by altered social interaction and restricted, repetitive behaviors or interests^2^. Autism is diagnosed about four times more in males than females, suggesting sex-specific factors may increase male vulnerability^3^. Previous work^4^ tested whether this bias is reflected in the genetic architecture of autism and other NDDs by comparing the enrichment of deleterious *de novo* coding variants in male and female NDD probands. Surprisingly, the most sex-biased genes showed an excess of *de novo* mutations in females, not males^4^. These genes—*DDX3X, HDAC8, WDR45, NAA10,* and *MECP2*—are all X-linked. Of these, only *MECP2* approached statistical significance when restricting diagnostic criteria to autism^4^. This likely reflects the fact that the equivalent loss-of-function coding mutations in hemizygous males produce even more severe phenotypes^5–9^ that are excluded from these NDD cohorts, whereas females are partially protected by mosaic expression from their second, wild-type allele. This raises the possibility that milder mutations in these X-linked genes—such as noncoding variants that alter expression levels—could produce a milder phenotype like autism, with a male-biased penetrance.

To investigate this, we focused on *MECP2,* a dosage-sensitive gene that encodes methyl CpG binding protein 2 (MeCP2). Strong loss-of-function variants in *MECP2* cause a profound disorder, Rett syndrome, in females^10^ but typically lead to severe neonatal encephalopathy and premature lethality in males^9^. Gain-of-function *MECP2* variants are often asymptomatic in females but cause profound neurodevelopmental phenotypes in males^11^, termed *MECP2* duplication syndrome (MDS). In both disorders, females are protected from more severe phenotypes by mosaic expression of wild-type *MECP2* on their second X chromosome. This holds true even for mild hypomorphic *MECP2* variants, such as p.A140V, which causes intellectual disability and psychiatric symptoms in males, while female carriers are healthy or minimally affected^12,13^. This led us to hypothesize that other mild *MECP2* variants, such as mutations in cis-regulatory elements (CREs), might cause non-syndromic neurological phenotypes like autism, particularly in males.

This hypothesis is supported by mouse models of the *Mecp2* allelic spectrum. While *Mecp2* null^14,15^ or over-expression alleles^16^ recapitulate Rett syndrome or MDS, respectively, mutations that lead to moderate changes in *Mecp2* expression levels can cause milder autism-like phenotypes. Deleting a conserved intronic enhancer (“CRE 2”) decreases *Mecp2* expression by 30%, causing social deficits, anxiety, and hyperactivity^17^. Conversely, deleting a conserved upstream repressor (“CRE 6”) increases gene expression by 50% and causes social defects, hypoactivity, and anxiety, and impairs motor function, learning, and memory^17^. This indicates that there is a narrow range for MeCP2 levels in healthy brain development, and that human variants in CREs that tip *MECP2* expression levels out of this range may cause autism in hemizygous males.

To begin testing this hypothesis, we took two parallel approaches: 1) test the functional impact of rare *MECP2* noncoding variants found in male autism probands, and 2) expand the catalog of *MECP2* CREs and improve their functional analysis to inform future variant discovery and interpretation. To this end, we performed massively parallel reporter assays (MPRAs) in induced pluripotent stem cell (iPSC)-derived neurons (iNeurons) to test autism-associated variants for altered CRE activity and to determine which regions of *MECP2* CREs contain active transcription factor (TF) binding sites (TFBSs). We identified multiple CRE sequences with clear activator or repressor binding sites and identified the most likely corresponding transcription factors (TFs). We also found autism-associated variants near or within these binding sites that impact CRE function. Finally, we validated a promoter variant identified in a male proband with non-syndromic autism and ADHD, discovering that it lowers endogenous *MECP2* expression by ∼30%. This study is the first to link noncoding *MECP2* variants to male-biased, non-syndromic autism, highlighting the potential for noncoding variants in other X-linked NDD genes to contribute to male-biased autism.

## Results

### A massively parallel reporter assay to dissect MECP2 cis-regulatory elements

To build a catalog of *MECP2* CREs in relevant human brain tissues, we examined regions of open chromatin from two independent human fetal prefrontal cortex datasets^18,19^ (Fig. 1a). We focused on the region between *OPN1LW* and *IRAK1*, as previous work has shown that a bacterial artificial chromosome encompassing this region is sufficient to reproduce the endogenous *MECP2* expression^16^, demonstrating that it contains all required CREs. As expected, we identified open chromatin at the *MECP2* promoter, and at the two previously identified^17^ regulatory elements, CRE 2 and CRE 6. We also found two unexplored regions of open chromatin, one in intron 2 close to CRE 2 (which we call “CRE A”) and one between the promoter and CRE 6 (“CRE B”). Unlike CREs 2 and 6, these elements are not conserved to mice, although they are conserved across multiple nonhuman primate species (Supplementary Fig. 1a-b).

**Figure 1.**
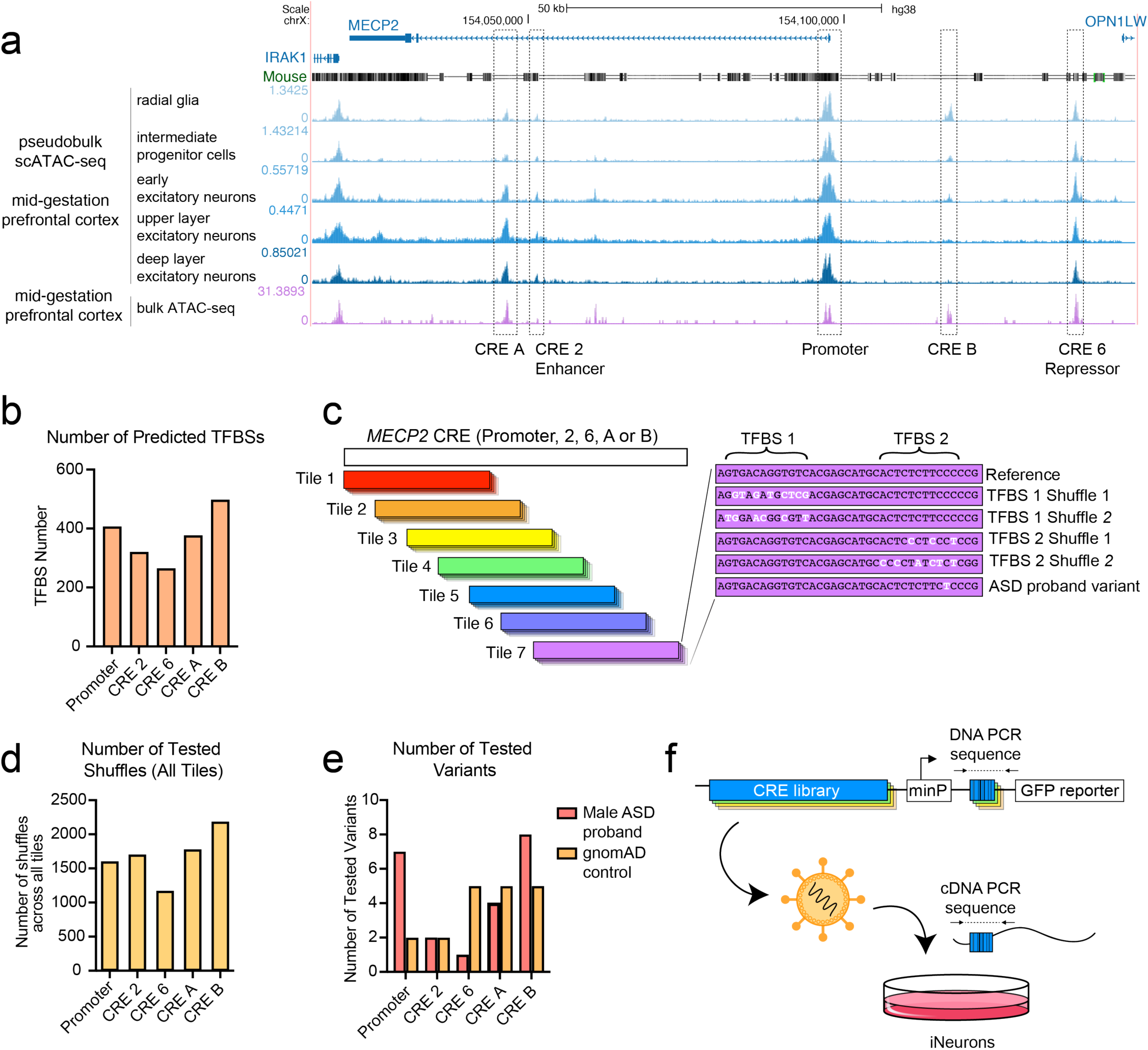
MPRA library design. **(a)** UCSC Genome Browser display of the *MECP2* locus with tracks showing conservation to mouse and two independent ATAC-seq datasets of mid-gestational prefrontal cortex^18,19^. **(b)** Number of predicted TFBSs within five *MECP2* regulatory elements. **(c)** Cartoon schematic of MPRA library design showing the tiling design and the nucleotide shuffling of predicted TFBSs. **(d)** Total number of TFBS mutations tested across all tiles, for each regulatory element. **(e)** Number of autism-associated variants and control variants tested across all tiles, for each CRE. **(f)** Cartoon of MPRA construct and experimental design.

CREs contain motifs that are recognized by TFs, which act in concert and with co-factors to direct gene expression^20^. We identified hundreds of potential TF motifs in each of the CREs (Fig. 1b). To test which ones actively contribute to CRE function, we designed a lentivirus MPRA to systematically perturb the motif and evaluate the effect on gene expression. As *MECP2* is most critical in neurons^15^, we performed this assay in human iNeurons, using *Ngn2* to drive differentiation into cortical glutamatergic neurons^21^. To test the whole CRE sequence within the constraints of an oligonucleotide library, we broke each CRE into overlapping 270 bp tiles, with a sliding window of 70 bp (Fig. 1c). In each tile, we predicted all possible TF motifs, then disrupted each motif by randomly shuffling its nucleotides, two independent times (Fig. 1c). Altogether, this introduced over 1,000 mutations in each CRE (Fig. 1d).

We also included a curated set of single-nucleotide variants identified in individuals with autism, derived from analyzing over 8,000 families across 14 different autism cohorts. We selected variants that were: 1) present in one of the five defined CREs; 2) ultra-rare (private to a single family); 3) found only in male probands; and 4) inherited from unaffected mothers. This resulted in 22 variants across the five CREs (Fig. 1e). As negative controls, we also included 19 gnomAD variants that were each found in at least 50 males assumed to be neurotypical (i.e., not designated as cases in neuropsychiatric studies) (Supplementary Table 1). As technical controls, we included two sets of sequences previously shown to have high or low activity in the iNeuron MPRA assay^22^.

We followed an established protocol for lentivirus MPRA^23^, performing four replicates in iNeurons (Fig. 1f). We achieved a median value of ∼135 barcodes per element (Supplementary Fig. 2a) and observed a strong correlation across the four replicates (Supplementary Fig. 2b). Our control sequences behaved as expected, with negative control sequences showing low log_2_(RNA/DNA) values and the positive control sequences with high log_2_(RNA/DNA) values (Supplementary Fig. 2c). Altogether, this satisfied our quality control criteria, indicating that our assay can reliably measure CRE activity in iNeurons.

### Annotating active TFBSs in MECP2 CREs

We first examined which shuffled TF motifs significantly impacted MPRA activity compared to their matched reference, then mapped these back to the full CRE sequence. Shuffled sequences and their effect sizes are provided in Supplementary Table 2. To visualize results, we plotted the average effect size of each shuffle mutation across the four replicates, sorting each mutation into 50 bp bins. Each motif is represented by multiple datapoints from independent shuffle mutations tested across multiple tiles. A motif that reproducibly contributes to CRE activity appears as a cluster of datapoints within a single bin.

We then defined “hotspot” regions with robust evidence of activator or repressor binding. Four out of five CREs (CRE 2, 6, A, and the proximal promoter) contained at least one hotspot (Supplementary Fig. 3a, Supplementary Table 3). Consistent with its positive regulatory role, the proximal promoter contained an activator-binding hotspot (Supplementary Fig. 3a). The known enhancer CRE 2 contained four activator-binding sites and three repressor-binding sites (Supplementary Fig. 3a, Fig. 2a), and the known repressor CRE 6 contained two strong repressor-binding sites and one activator-binding site (Supplementary Fig. 3a-b). These data align with previous findings that both activator and repressor motifs can work together to fine-tune a CRE’s regulatory output^24^. Notably, the CTCF binding site previously identified in CRE 6^17^ does not reach statistical significance here yet has a subtle repressive effect (Supplementary Fig. 3b-c), suggesting that it retains some repressor function even without its endogenous chromatin context. Finally, we evaluated the two newly described CREs. One element, CRE B, had no active TFBSs (Supplementary Fig. 3a, Supplementary Fig. 3d). This is consistent with reduced chromatin accessibility at CRE B in differentiated neurons compared to radial glia (Fig. 1a) and suggests that CRE B may regulate *MECP2* in neural stem cells rather than neurons. CRE A, in contrast, contained three repressor-binding sites and no activator-binding sites (Supplementary Fig. 3a), consistent with a role as a repressor in neurons.

**Figure 2.**
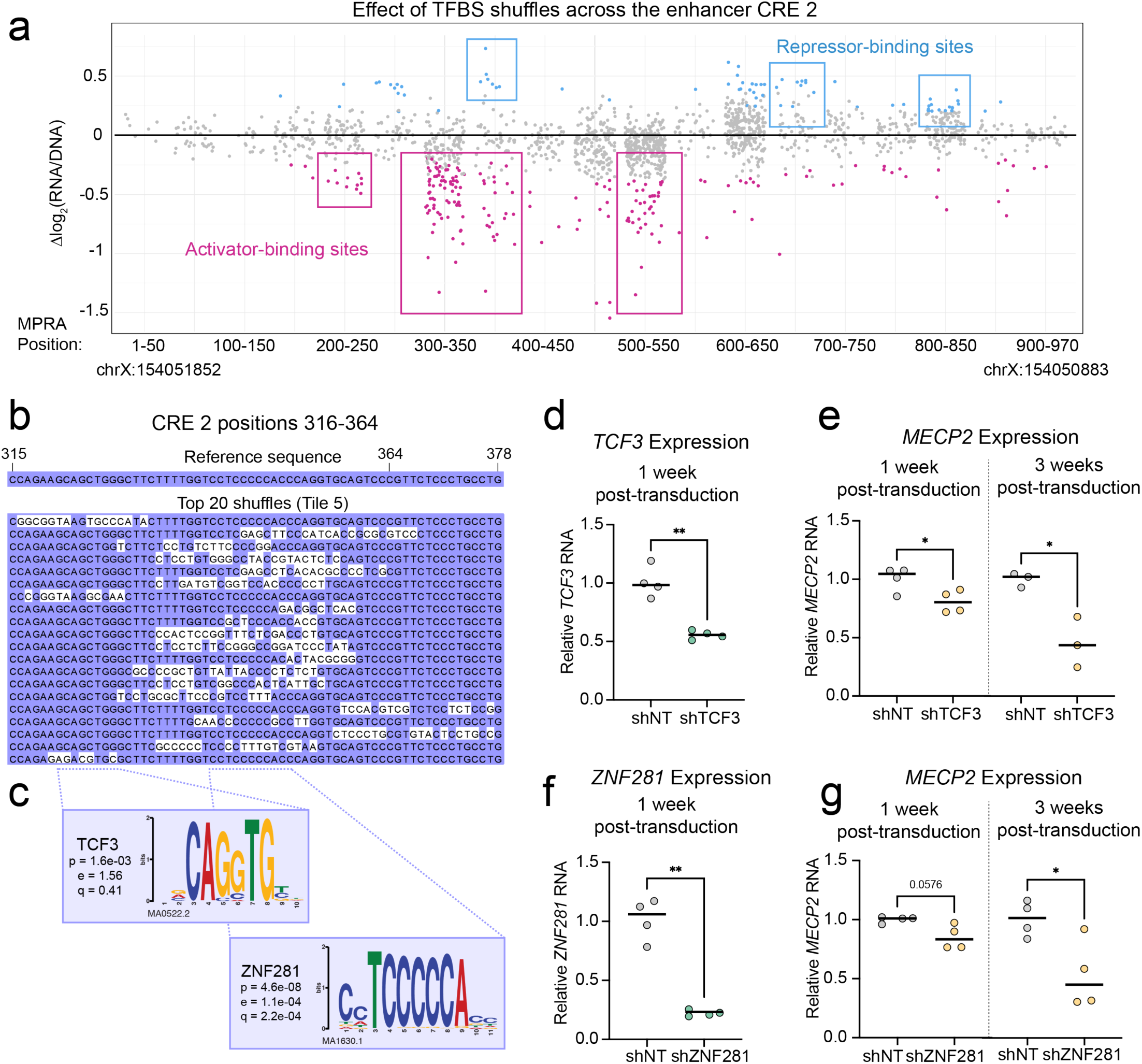
Identifying active TFBSs in the CRE 2 enhancer. **(a)** Effect of shuffling TFBSs on CRE 2 activity. Each datapoint represents a shuffle mutation, with statistically significant (p_adj_<0.05) values represented in blue (increased expression) or pink (decreased expression). Data are shown in 50 bp bins across the CRE, with starting and ending hg38 coordinates for reference. Repressor- or activator-binding sites are marked in blue or pink boxes, respectively. **(b)** Alignment of sequences underlying one activator-binding site, with the reference sequence above. Each line shows an independently tested sequence. Nucleotides matching the reference are displayed with a purple background, while mutations are displayed with a white background. **(c)** Position weight matrices (PWMs) for TCF3 and ZNF281, two TFs with best motif matches to the disrupted sequences. The p value reports the statistical significance in similarity between motif and sequence, while the e and q values report the expected number of false positives from the match process and the false discovery rate, respectively^25^. **(d)** RT-qPCR of *TCF3* cDNA one week after transduction with shRNAs targeting *TCF3* RNA, relative to a non-targeting (NT) control. **(e)** RT-qPCR of *MECP2* cDNA one week (left) and three weeks (right) after transduction with shRNAs targeting *TCF3* RNA, relative to a non-targeting (NT) control. **(f)** RT-qPCR of *ZNF281* cDNA one week after transduction with shRNA targeting *ZNF281* RNA, relative to a non-targeting (NT) control. **(g)** RT-qPCR of *MECP2* cDNA one week (left) and three weeks (right) after shRNA transduction targeting *ZNF281* RNA, relative to a non-targeting (NT) control. For **(e-g)**, statistical significance was calculated with a Welch’s t-test; * p<0.05, ** p<0.001. Each timepoint represents an independent iNeuron differentiation.

We then extracted the sequences underlying the TF binding hotspots and aligned them to assess whether the mutations overlapped with each other and to predict which motifs were driving the difference in activity. For example, in CRE 2, we focused on an activator-binding site with a large effect between positions 300 and 400 (Fig. 2a), where we found the critical sequence between positions 316-364 (Fig. 2b). We then used an unbiased, orthogonal approach to independently test which motifs contributed the most to the change in activity. Briefly, we identified nucleotide sequences (i.e., k-mers) with the greatest effect on CRE activity, matched each to known motifs using TomTom^25^, and refined iteratively to the smallest k-mer that optimized the motif match. We then prioritized motifs corresponding to TFs expressed in iNeurons^26,27^ and the developing brain^28^ to focus on biologically relevant candidates. This approach improved the mapping of TFBSs with strong contribution to CRE activity, refining the list of candidate TFs that may mediate its regulatory activity. For the CRE 2 activator at position 316-364, we identified an E-box motif and zinc finger motif with strong matches to TCF3 and ZNF281 (Fig. 2c). Lentiviral shRNA knockdown of *TCF3* in iNeurons (Fig. 2d) reduced *MECP2* expression by 20% after one week and by ∼55% two weeks later (Fig. 2e). Similarly, *ZNF281* knockdown reduced *MECP2* RNA by ∼15% (p=0.0576) after one week and by 47% two weeks later (Fig. 2f-g). These results support a role for TCF3 and ZNF281 as transcriptional activators of *MECP2*. They also demonstrate that mapping the most dynamic *cis*-acting sequences within a CRE can predict relevant upstream *trans*-acting factors. A complete list of k-mers and associated motifs that impact activity is available in Supplementary Table 4.

### CRE 2 TFBS activity is influenced by flanking sequence context

The MPRA’s tiling design allowed us to investigate whether the activity of these TFBSs is impacted by the context of the surrounding nucleotides. To test this, we focused on the strong activator-binding sites we detected in CRE 2. The shuffled sequences at the TCF3/ZNF281 motifs were tested with four different flanking sequences, moving from the far 3’ end of the tile (Tile 2) to the far 5’ end of the tile (Tile 5) (Fig. 3a). Overall, the mutations that decreased MPRA activity in one tile similarly decreased activity in the other tiles, indicating that this TFBS is active regardless of the flanking sequence (Fig. 3b). However, we discovered the opposite effect at the nearby activator-binding site at position 500-550 (Fig. 2a). Through sequence alignment and our motif prediction strategy, we narrowed the key region to the poly-adenine sequence within position 510-548 (Fig. 3c), with the best motif prediction matching a ZNF362 motif (Supplementary Table 4). Here, the mutations that decreased MPRA activity in Tile 5 lost their effect when the same mutation was tested in Tiles 6, 7, or 8 (Fig. 3d). One possibility is that moving from Tile 5 to Tile 6 adds a 3’ repressor-binding site that neutralizes the upstream ZNF362 activator site. However, the nearest repressor-binding site at 650-700 (Fig. 2a) is not included in the sequence until Tile 7 (Supplementary Table 2) and thus cannot explain the loss of activator effect in Tile 6. The other possibility is that moving from Tile 5 to Tile 6 loses a 5’ activating partner. Indeed, the TCF3/ZNF281 site is included in Tile 5 but mostly excluded from Tile 6 (Fig. 3a). This suggests that the factor(s) binding to the ZNF362 motif rely on a partnership with those binding to the TCF3/ZNF281 motifs to drive CRE activity.

**Figure 3.**
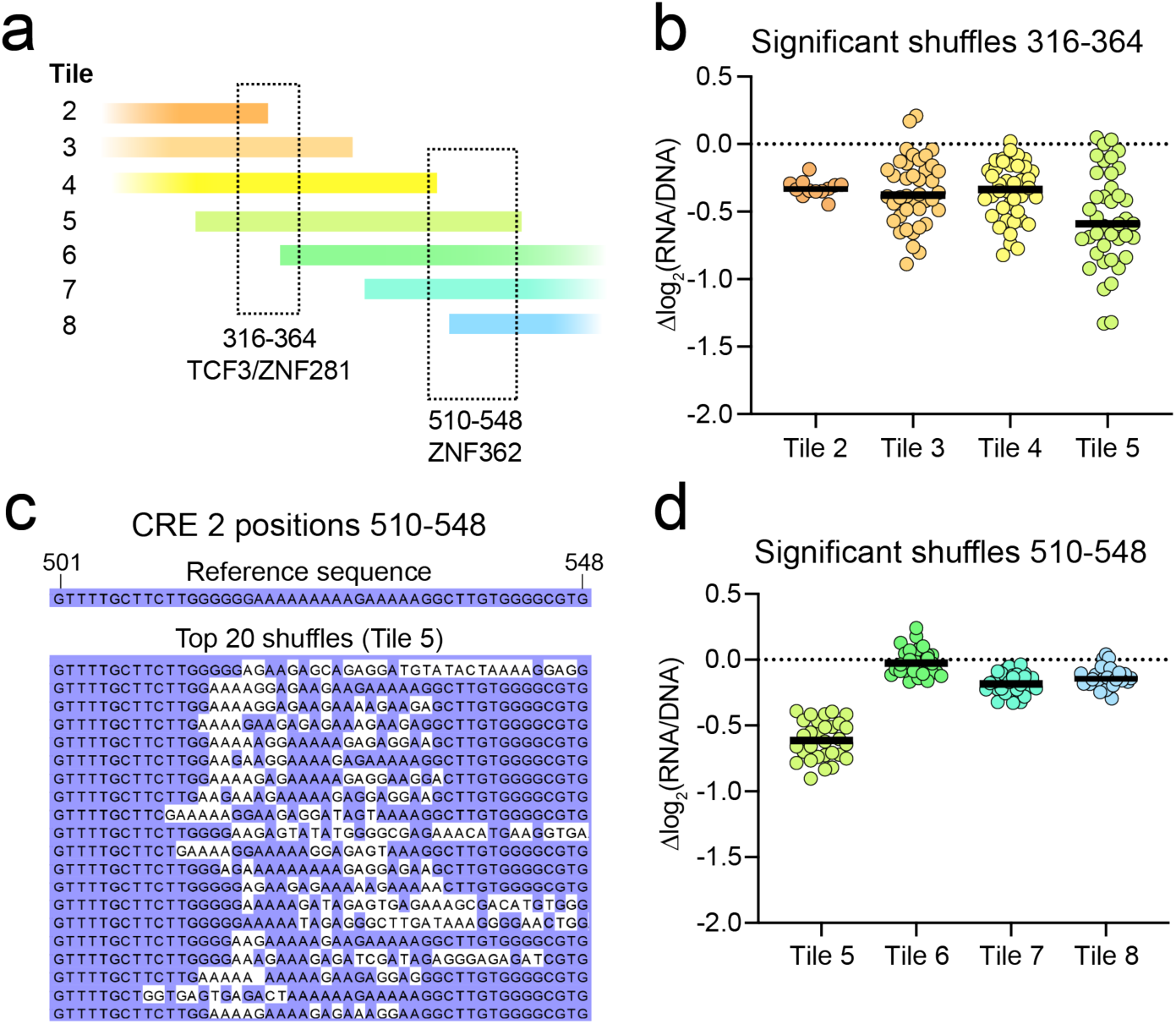
MPRA detects context-dependent TF binding activity. **(a)** Cartoon diagram showing two active TF binding sites in CRE 2 and the tiles including their sequence. **(b)** Effect size of the statistically significant (p_adj_<0.05) shuffles in positions 316-364, shown for each tile that includes the mutation. **(c)** Alignment of sequences underlying the activator binding site at position 510-548, with the reference sequence above. Each line shows an independently tested sequence that decreases activity in Tile 5. Nucleotides matching the reference are displayed with a purple background, while mutations are displayed with a white background. **(d)** Effect size of the significant shuffles in positions 510-548, shown for each tile that includes the mutation.

### A male-biased autism variant in the novel CRE A repressor results in a loss of repressor activity

To expand our study beyond the known CRE 2 and CRE 6 elements, we turned to the primate-specific element CRE A. We identified multiple hotspot regions with repressor-binding activity, including a strong hotspot at positions 819-894 (Fig. 4a). This signal was driven by a zinc finger motif and a T-box motif, matching ZNF282 and MGA motifs (Fig. 4b, Supplementary Table 4). We also tested the effects of four maternally inherited, ultra-rare CRE A variants found in male autism probands. To ensure that the assay was well-calibrated, we included five variants from gnomAD as controls. One gnomAD variant, chrX:154046641:C:T, modestly but consistently increased CRE activity (Fig. 4c), suggesting that this degree of variation in CRE A activity is tolerated by unaffected individuals. Two of the four autism variants (−6746:A:C and −6128:G:A) significantly altered activity in one tile, but with an average effect size smaller than that of the gnomAD control variant. A third autism variant (−6121:G:A) did not significantly change activity in any test.

**Figure 4.**
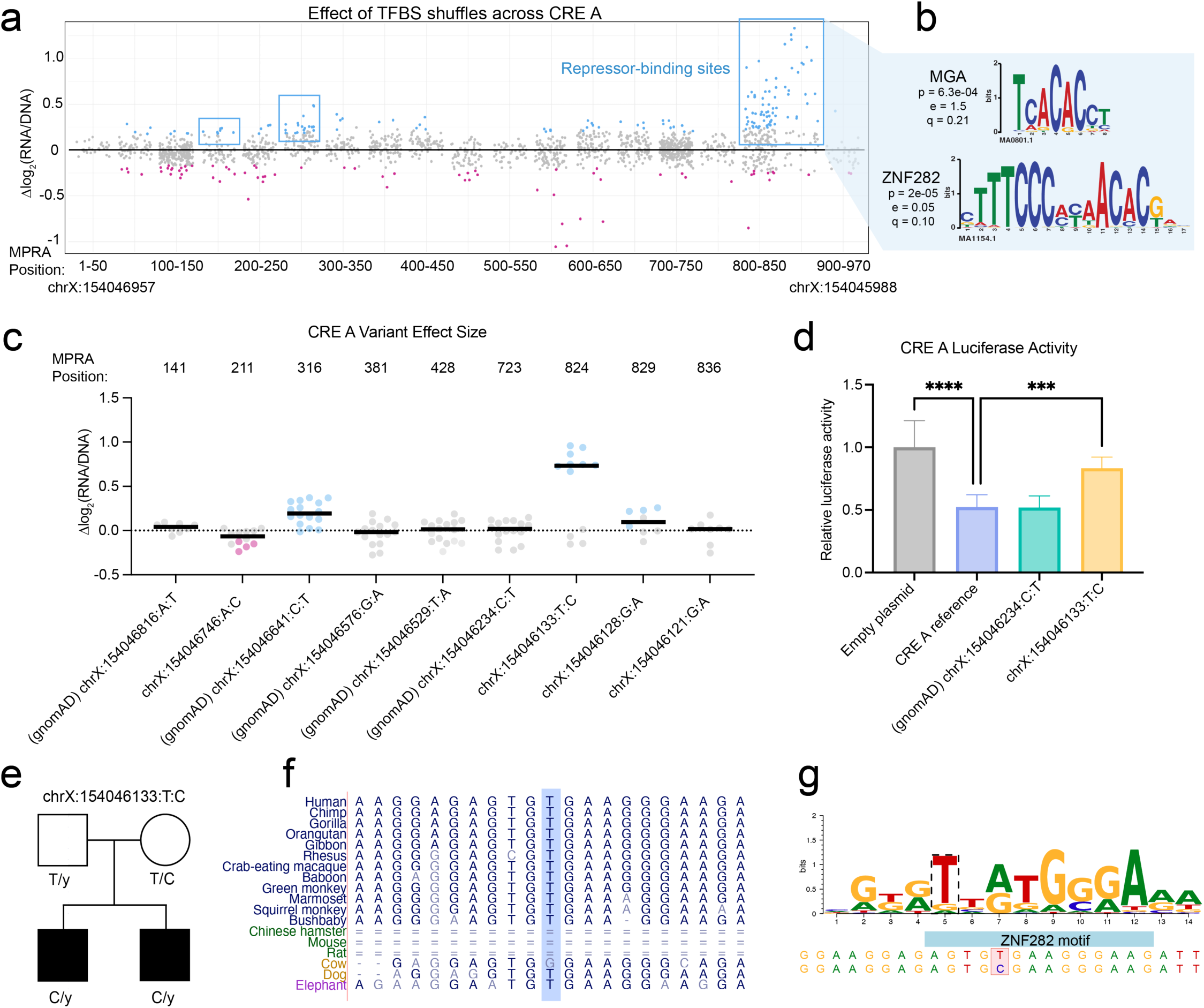
An autism-associated CRE A variant disrupts a repressor-binding site. **(a)** Effect of shuffling CRE A TFBSs. Each datapoint represents a shuffle mutation, with statistically significant (p_adj_<0.05) points represented in blue or pink. Data are shown in 50 bp bins across the CRE, with starting and ending hg38 coordinates for reference. Repressor binding sites are marked in blue boxes. **(b)** PWMs for MGA and ZNF282, two TFs with best motif matches to the disrupted sequences. The p value reports the statistical significance in similarity between motif and sequence, while the e and q values report the expected number of false positives from the match and the false discovery rate, respectively^25^. **(c)** Effect of autism and gnomAD control variants on MPRA activity. The X axis displays the hg38 coordinate, reference allele, variant allele, with the position within the MPRA listed on top. Each datapoint shows the variant’s effect size within a given tile. Statistically significant (p_adj_<0.05) values are shown in pink or blue for decreased or increased activity, respectively. **(d)** Luciferase assay comparing the activity of the CRE A reference sequence to an empty plasmid, to a gnomAD control variant, and to the autism variant chrX:154046133:T:C. Statistical significance was calculated using a one-way ANOVA with Dunnett’s multiple comparisons test; **** p<0.0001, *** p<0.001. Data is representative of three independent trials across three differentiations. **(e)** X-linked segregation of the variant with an autism phenotype in family KP5115 (Pelphrey cohort). **(f)** UCSC Browser conservation tracks for the chrX:154046133 nucleotide. **(g)** PWM for the motif predicted to be disrupted by chrX:154046133:T:C, matching the reverse complement of the motif in **(b)**.

The other autism variant (chrX:154046133:T:C) significantly increased activity beyond the effect of the benign gnomAD control (Fig. 4c). Notably, it is located at MPRA position 824, within the strong repressor hotspot located at position 819-894. To validate this, we performed a luciferase reporter assay with the entire 970 bp CRE A region cloned upstream of a minimal promoter. We found that CRE A repressed luciferase activity below that of an empty vector, consistent with the strong repressor-binding activity we observed in the MPRA (Fig. 4d). While a nearby gnomAD variant resembled the reference sequence, the variant chrX:154046133:T:C significantly increased activity (Fig. 4d). To assess the genetic evidence for pathogenicity, we evaluated the associated pedigree. The variant was identified in a pedigree with two affected male probands, who both inherited the C allele from their unaffected mother (Fig. 4e). Neither proband carried any *de novo* coding variants or copy number variants (CNVs) in known autism genes that might otherwise explain their phenotype, although both harbored inherited missense variants of uncertain significance in NDD genes (Supplementary Table 5). The chrX:154046133 position and surrounding nucleotides are conserved among primates and other large mammals (Fig. 4f), indicating this region may be important for *MECP2* expression. Finally, we used motifbreakR^29^ to predict whether this variant disrupts a TFBS. We identified the same ZNF282 motif (Fig. 4g) that we found in our independent motif shuffle analysis (Fig. 4b). Altogether, the data support the potential pathogenicity of this variant, laying a foundation for implicating a novel *MECP2* CRE in male-biased autism.

### A male proband variant in the proximal promoter impacts MECP2 expression

We next investigated the architecture of the *MECP2* proximal promoter, selecting a 970 bp sequence spanning 1021-51 bp upstream of the transcription start site (TSS). This excludes the GC-rich core promoter that binds the transcription initiation machinery^30^. Across this ∼1 kb sequence, we identified a single activator-binding hotspot at MPRA coordinates 918-955 (Fig. 5a) corresponding to a SP/KLF motif and an E-box motif. Motif matching identified KLF7 and ZEB1 as the strongest candidates with robust expression in iNeurons (Fig. 5b, Supplementary Table 4).

**Figure 5.**
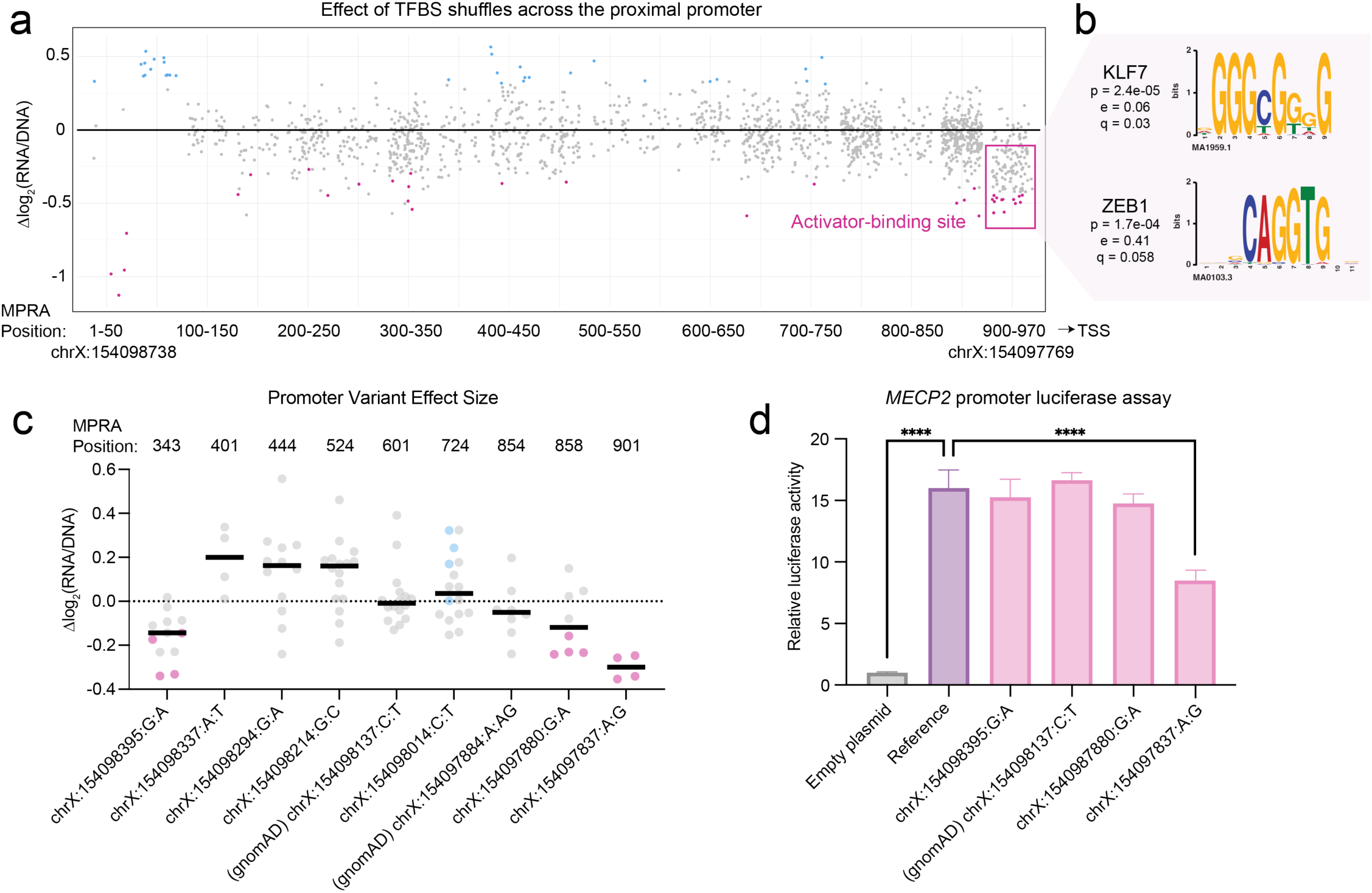
The 3’ end of the *MECP2* proximal promoter drives activity. **(a)** The effect of shuffling TFBSs in the proximal promoter. Statistically significant (p_adj_<0.05) values are shown in blue or pink. The pink box denotes an activator binding site. **(b)** PWMs for KLF7 and ZEB1. The p value reports the statistical significance in similarity between the motif and the disrupted sequence, the e value reports the expected number of false positives from the matching process, and the q value reports the false discovery rate^25^. **(c)** Effect of autism and gnomAD control variants on MPRA activity. The X axis displays the hg38 coordinate, reference allele, and variant allele, with the position within the MPRA shown on top. Each datapoint shows the variant’s effect size within a given tile. Statistically significant (p_adj_<0.05) values are shown in pink or blue for decreased or increased activity, respectively. **(d)** Luciferase assay comparing the activity of the proximal promoter reference sequence to an empty plasmid, to a gnomAD control variant, and to three variants identified in autism probands. Statistical significance was calculated using a one-way ANOVA with Dunnett’s multiple comparisons test; **** p<0.0001. Data is representative of three independent trials across three differentiations.

We also tested seven maternally inherited variants from male autism probands, alongside two gnomAD variants. These controls did not affect regulatory activity (Fig. 5c). Four of the seven autism proband variants significantly changed regulatory activity in one tile, and three of these (−8395:G:A, −7880:G:A, and −7837:A:G) had a larger average impact than either gnomAD variant (Fig. 5c). However, they did not change activity when tested in additional tiles, indicating that these effects may be context-dependent or false positives. (Variant −7837:A:G was positioned so close to the 3’ end of the design that it could only be tested in one tile.) To clarify these data, we turned to the luciferase reporter assay, using the entire 970 bp sequence upstream of a minimal promoter. As expected, the reference sequence drove strong luciferase activity compared to an empty plasmid control (Fig. 5d). Two of the three autism variants did not change this activity (Fig. 5d). However, the variant −7837:A:G significantly decreased luciferase activity (Fig. 5d). This variant is located at MPRA position 901, 17 bp away from the KLF7/ZEB1 active motif and 120 bp upstream of the annotated TSS (Supplementary Fig. 4).

To further investigate the pathogenicity of this variant, we took a closer look at the genetic and phenotypic data of the proband. This variant was identified in a pedigree from the Simons Simplex Collection (SSC)^31^ and was passed from an unaffected mother to her affected son, but not her neurotypical son (Fig. 6a). We did not identify any *de novo* coding variants or CNVs in known autism genes that might explain the affected son’s phenotype. Although the proband did inherit a missense variant of uncertain significance in *NAA15*, this was also transmitted to his unaffected sibling (Supplementary Table 5). We were surprised to find a deleterious variant so close to the core *MECP2* promoter in a male case of non-syndromic autism, and so we carefully evaluated the extensive phenotypic data collected by SSC to rule out Rett syndrome features. The proband was diagnosed with autism and ADHD but had a full-scale IQ of 97 and normal verbal skills, with no motor deficits or seizures (Fig. 6b)—consistent with non-syndromic autism.

**Figure 6.**
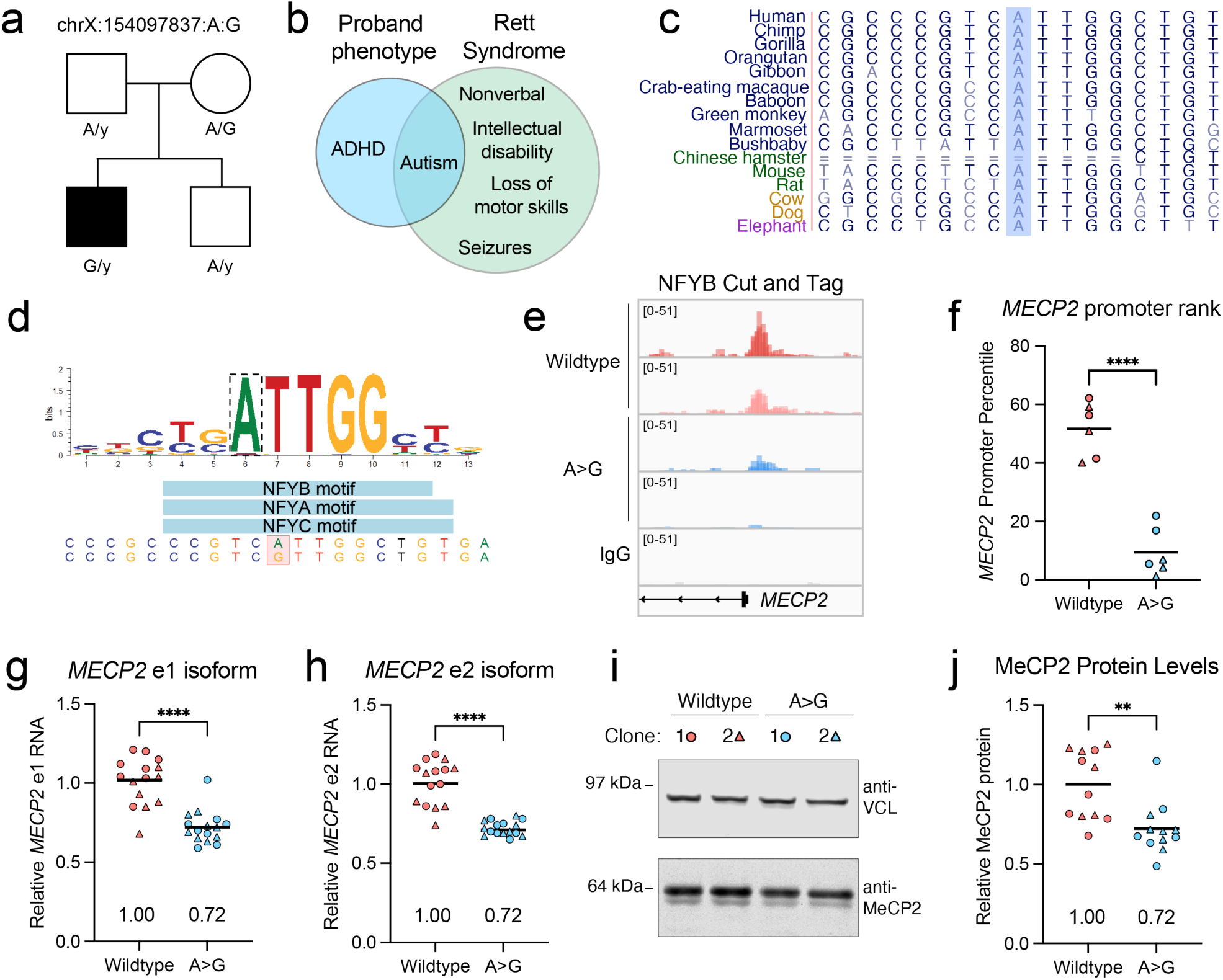
An autism-associated promoter variant reduces *MECP2* expression. **(a)** X-linked segregation of the variant with an autism phenotype in family SSC 14435. **(b)** Comparison of the proband phenotype with typical features of Rett syndrome. **(c)** UCSC Browser conservation tracks for the chrX:154097837 nucleotide. **(d)** PWM for the motif predicted to be disrupted by −7837:A:G. **(e)** NFY-B CUT&Tag signal at the *MECP2* promoter. For NFY-B, each track shows the overlay of three technical replicates. For IgG, the overlay of all twelve tracks is shown. **(f**) *MECP2* promoter rank across all promoters bound by NFY-B in wild-type and A>G iNeurons. (**g-h)** RT-qPCR for the *MECP2* e1 **(g)** and e2 **(h)** isoforms in iNeurons. Relative expression was calculated using the 2^−ΔΔCT^ method, with GAPDH used as a normalization control. **(i)** Representative western blot image and quantification **(j)** for unedited and A>G iNeurons. MeCP2 values are normalized to VCL. For (**f-j**), circles and triangles represent two independently derived clones for each genotype. Statistical significance was calculated with a Welch’s t-test; ** p< 0.01, *** p<0.001, **** p<0.0001. For **(g-j)**, data is representative of at least two independent differentiations.

The −7837:A:G variant affects a highly conserved residue in the proximal promoter (Fig. 6c) and is predicted to disrupt a NFY binding site (Fig. 6d). NFY is a trimeric complex (with -A, -B, and -C subunits) involved in nucleosome depletion and transcription initiation at promoters^32,33^ and regulates both housekeeping^34^ and neuron-specific genes^35^. Intriguingly, NFY-B binds to the *MECP2* promoter in ChIP-Seq datasets from iPSCs, K562 cells, and GM12878 cells (Supplementary Fig. 5a), indicating that this complex may regulate *MECP2* expression. To test this, we used CRISPR-Cas9 to edit this variant into the native locus in iPSCs. After editing, we isolated clonal lines with or without the variant, using matched controls that were subject to nucleofection and single-cell cloning but did not undergo editing. We selected two variant clones and two wild-type clones to differentiate into iNeurons, collecting cells after 4 weeks of neuronal differentiation. To determine whether the −7837:A:G variant impairs NFY binding, we performed CUT&Tag for NFY-B across our four clones, as well as for IgG as a negative control. NFY-B iNeuron peaks substantially overlapped with NFY-B peaks in the iPSC, K562, and GM12878 datasets, with ∼35% of peaks shared between all four cell types, and another 15% shared between iNeurons and 1-2 other cell types (Supplementary Fig. 5b). The remaining peaks may be neuron-specific or protocol-dependent (i.e., CUT&Tag in this study compared to ChIP-Seq). As expected, the iNeuron peaks were strongly enriched for promoters (Supplementary Fig. 5c), and *de novo* motif analysis identified enrichment of the NFY motif ATTGG (Supplementary Fig. 5d).

After confirming dataset quality, we next evaluated NFY-B signal over the *MECP2* promoter, observing a notable decrease (Fig. 6e). To quantify this relative to the broader promoter-associated NFY-B binding, we ranked the NFY-B signal at all promoter peaks within each replicate. We then calculated the percentile for the *MECP2* promoter for each replicate. While the *MECP2* promoter is in the 51^st^ percentile of peaks in the wild-type samples, it falls to the 9^th^ percentile in the −7837:A:G samples (Fig. 6f). This demonstrates that the −7837:A:G variant significantly impairs NFY binding.

To assess the impact on *MECP2* gene expression, we harvested RNA and protein from our four-week-old iNeurons. Using qRT-PCR, we detected a ∼30% decrease in both known isoforms of *MECP2*, e1 and e2 (Fig. 6g-h). Consistent with these findings, Western blot analysis showed that MeCP2 protein level was also decreased by ∼30% (Fig. 6i-j). Altogether, the data support the pathogenicity of the chrX:154097837:A:G variant and support the hypothesis that hypomorphic, noncoding *MECP2* variants cause male-biased non-syndromic autism.

## Discussion

We explored the hypothesis that noncoding variants impacting dosage-sensitive, X-linked NDD genes contribute to male-biased autism, focusing on *MECP2* as initial proof-of-concept. We performed a MPRA in iNeurons to define critical TFBSs in *MECP2* CREs, predict their trans-acting partners, and test the functional consequences of ultra-rare variants identified in male autism probands. We mapped key activator- and/or repressor-binding sites for four of the five CREs we interrogated. We also identified two autism-associated variants that impact CRE function, one in the proximal promoter and one in the newly described CRE A repressor. Consistent with the expected male sex bias, both variants were inherited from an unaffected mother. Modeling the promoter variant at the endogenous *MECP2* locus in iNeurons revealed that it reduced RNA and protein by ∼30%, a degree that is sufficient to produce autism-like phenotypes in mice^17^.

One of the goals of this study was to develop a strategy for improving the annotation of key active sequences within a CRE, to narrow the search space for noncoding pathogenic variants and help prioritize variants for follow-up studies. While not all proband variants overlapping an active TFBS affected CRE activity, the two variants that altered activity in our assay were within or close to a strong TFBS. While the CRE A variant perfectly aligned with the major CRE A repressor-binding site, the promoter variant was 17 bp away from the strongest activator-binding sites. This may be explained by our focus on non-syndromic autism cohorts; a variant directly impacting the activator motifs might cause more features of Rett syndrome or even premature lethality in males. Overall, the orthogonal dataset for TFBSs helped bolster our interpretation of the human proband variants. We also mitigated false positives by focusing on variants that impacted function in more than one MPRA tile, and by including variants from neurotypical gnomAD controls to define a threshold for a benign change in reporter expression. These careful design considerations may help future MPRA studies prioritize the most likely pathogenic variants for additional mechanistic studies.

We also predicted candidate TF motifs underlying each of these activator- and repressor-binding sites, matched them to TFs expressed in iNeurons, and demonstrated a functional relationship between selected TFs and *MECP2*. However, we are limited in our ability to comprehensively catalog which TFs (or which combination of TFs) directly regulate *MECP2* expression. This is partly because a motif can be matched with many different TFs that have similar position weight matrices, especially those within the same TF family^36^. Despite this, some of the TFs we identified are compelling candidates that have been implicated in NDDs, such as *KLF7*^37^. Others, such as *ZNF281* and *ZNF362*, have not been associated with disease but are highly constrained^38^ and are predicted to cause NDD^39^. This study opens the door for more detailed investigation of whether variants in these TFs cause NDD and whether altered *MECP2* expression contributes to the pathophysiology.

We must also acknowledge the limitations of MPRAs, which take a relatively short sequence (in this case ∼300 bp) out of its genomic context to test its regulatory potential. Although we attempted to mitigate this with our tiling design and subsequent luciferase validations, we may have missed discovering TFBSs dependent on longer-range interactions within the CRE. Additionally, by cloning the library directly upstream of a minimal promoter, we cannot accurately model complex or distal interactions with the *MECP2* core promoter that arise from larger chromatin loops. This limitation is illustrated in the case of the CRE 6 CTCF site, which appears to act as an insulator at the endogenous locus^17^ yet is only weakly repressive in the MPRA context.

In summary, this work refines the cis-regulatory map of *MECP2*, expands its allelic series to include noncoding variants linked to non-syndromic autism in males, and establishes a generalizable framework for interrogating cis-regulatory variation in X-linked NDD genes that may underlie a fraction of autism’s missing heritability and male sex bias.

## Acknowledgements

We are grateful for the whole genome sequencing data available from ANVIL and SFARIBASE from participating autism families including the Simons Simplex Collection (SSC) and associated principal investigators (A. Beaudet, R. Bernier, J. Constantino, E. Cook, E. Fombonne, D. Geschwind, R. Goin-Kochel, E. Hanson, D. Grice, A. Klin, D. Ledbetter, C. Lord, C. Martin, D. Martin, R. Maxim, J. Miles, O. Ousley, K. Pelphrey, B. Peterson, J. Piggot, C. Saulnier, M. State, W. Stone, J. Sutcliffe, C. Walsh, Z. Warren, E. Wijsman). We also thank the staff of the Human Stem Cell and Neuronal Differentiation Core at Baylor College of Medicine and Dr. Aleksander Bajic for his expert assistance. We are grateful to Dr. Nadav Ahituv and Dr. Chengyu Deng for their guidance on performing lentiMPRA in iNeurons, as well as Dr. Li Gan for sharing the WTC11-Ngn2 iPSC line.

## Funding statement

This work was supported by grants from the Simons Foundation (SFARI #SFI-AN-AR-Sex Differences-00017956-01 to A.C., −02 to E.E.E., −03 to H.Y.Z., and −04 to T.N.) Research was also supported by the National Institutes of Health (NIH) under Award Number F32HD116501 to R.M-S and R01MH101221 to E.E.E. The use of the Baylor College of Medicine Human Stem Cell and Neuronal Differentiation Core facility and equipment was supported by the NIH under Award P50HD103555, NIH P30CA125123, and S10OD028591. The Duncan NRI Bioinformatics Core is supported by the Chan Zuckerberg Initiative (2023-332162), the Eunice Kennedy Shriver National Institute of Child Health and Human Development of the NIH (under Award P50HD103555), the Chao Endowment, the Huffington Foundation, and the Jan and Dan Duncan Neurological Research Institute at Texas Children’s Hospital. The content is solely the responsibility of the authors and does not necessarily represent the official views of the NIH. T.W. is supported by the Brain Science and Brain-like Intelligence Technology-National Science and Technology Major Project (2025ZD0218000), and the National Natural Science Foundation of China (82471194).

E.E.E. and H.Y.Z. are investigators of the Howard Hughes Medical Institute. This article is subject to HHMI’s Immediate Access to Research policy, which requires that this article be made publicly available as initial and revised preprints deposited on a designated preprint server under a CC BY 4.0 license.

## Methods

### Cohorts and variant discovery

Short-read whole-genome sequencing (WGS) data with an average read depth of 30× for 30,916 individuals from the CCDG (Centers for Common Disease Genomics, available on anvilproject.org) autism cohorts were analyzed. In total, there are 8,245 families with autism, including 9,437 probands (1,917 females and 7,520 males) and 4,989 unaffected siblings (2,543 females and 2,446 males). Single-nucleotide variants (SNVs) and small insertions/deletions (indels; <50 bp) were identified using FreeBayes (v1.1.0-3-g961e5f3)^40^ and GATK (v3.7.0)^41^, where the intersection set of variants were used for downstream analyses. Privately inherited variants (PIVs) on the X chromosome within the target regions (chrX:154,097,769-154,098,738, chrX:154,050,883-154,051,852, chrX:154,136,537-154,137,226, chrX:154,116,410-154,117,379, and chrX:154,045,988-154,046,957) were identified following the approach described previously^42^ with minor modifications. Briefly, we traced Mendelian inheritance for variants supported by both GATK and FreeBayes (GRCh38) within the target regions and calculated the frequency of heterozygous variants in parents. For candidate variants only observed in a single family, specifically those transmitted to male probands from unaffected mothers, we applied additional filtering criteria (QUAL > 50) and per-sample sequencing depth (DP ≥ 20). In total, we identified 22 PIVs corresponding to an allele frequency of approximately 6.1 × 10⁻⁵ based on 16,490 unrelated parents. For the two families prioritized for follow-up studies (Pelphrey cohort family KP5115 and SSC family 14435), we evaluated PIVs and *de novo* mutations genome-wide to assess whether alternative variants may be responsible for the phenotype (Supplementary Table 5). Variant effects were annotated and predicted using ENSEMBL Variant Effect Predictor and the RefSeq transcript.

### MPRA design

Candidate CREs were selected from fetal brain ATAC-seq datasets^18,19^. The sequence under each peak was extended to 970 bp (CREs A, B, 2, and promoter) or 690 bp (CRE 6). Each CRE was then divided into 270 bp tiles offset by 70 bp, with a 200 bp overlap. To design the motif shuffling part of the library, TF motifs within each tile were identified using the MEME Suite tool FIMO^43^, using the CIS-BP v2.0 motif library. Each motif sequence was extracted, shuffled two independent times using the universalmotif package (shuffle sequence function, linear shuffling, k=1), then placed back into the tile. To build the human variant part of the library, the 22 PIVs were manually inserted into the relevant library tiles. As controls, variants from gnomAD identified in at least 50 males from the v3.1.2 non-neuro subset were included. Finally, positive and negative controls (100 sequences each) known to have high or low activity in iNeuron lentiMPRA were included^44^.

### LentiMPRA cloning

The lentiMPRA library was constructed as previously described^23^. Briefly, a 300 bp TWIST oligo pool (270 bp insert + two 15 bp PCR handles) was amplified by a 5-cycle PCR using NEBNext High-Fidelity 2x PCR Master Mix (NEB M0541L), with primers 5BC-AG-f01 and 5BC-AG-r01. This step added the minimal promoter and vector overhang sequences. The PCR product was purified using 1X HighPrep PCR magnetic beads (Magbio Genomics AC-60050). A second, 14-cycle PCR using NEBNext High-Fidelity 2x PCR Master Mix (NEB M0541L) was performed to add a 15 bp random barcode and vector overhang sequence downstream of the insert library (with primers 5BC-AG-f02 and 5BC-AG-r02). The amplicon was purified using gel extraction with NEB Monarch DNA Gel Extraction Kit (T1020S). The amplicon was then cloned into the vector pLS-SceI (Addgene 137725), prepared by linearizing with *Age*I-HF (NEB R3552S) and *Sbf*I-HF (NEB R3642S). The recombination reaction was performed using NEBuilder HiFi DNA Assembly Master Mix (NEB E2621L) at 50°C for 60 minutes. To remove any non-recombined vector, the reaction was digested with I-SceI (NEB R0694S) before purification with 1.8 volume of HighPrep PCR magnetic beads (Magbio Genomics AC-60050). The cleaned product was electroporated into 10-beta electrocompetent cells (NEB C3020K) using a MicroPulser Electroporator (Biorad 1652100) with the E.coli setting (1.8 kVolt). To recover, cells were incubated in outgrowth media (NEB C3020K) at 37°C for 1 hour with agitation, then plated on 15cm plates alongside 100mg/mL carbenicillin. Approximately 2.2 million colonies were collected from four plates, pooled, and midiprepped (Takara Bio 740420.50) to obtain an average of 200 barcodes per insert. Then, each barcode was associated with an insert sequence using Illumina sequencing. Illumina flow cell adapters were added with PCR NEBNext High-Fidelity 2x PCR Master Mix (NEB M0541L) and the primers pLSmP-ass-i741 and P7pLSmP-ass-gfp. The amplicon was gel-purified using NEB Monarch DNA Gel Extraction Kit (T1020S), then further cleaned with 1.8 volumes of HighPrep PCR magnetic beads (Magbio Genomics AC-60050). The library was sequenced on an Illumina MiSeq with 2×250 bp configuration to ∼22 million reads with custom primers (Read 1: pLSmP-ass-seq-R1, Read 2: pLSmP-ass-seqR2, Index read 1: pLSmP-ass-seq-ind1).

### iPSC culture

This study used the WTC11 iPSC cell line with a doxycycline-inducible Ngn2 integrated at the AAVS1 safe-harbor locus, generated by Dr. Li Gan’s laboratory^21^. iPSCs were cultured under feeder-free conditions using Matrigel coating (Corning 356231) and mTESR Plus media (Stemcell Technologies 100-0276). iPSCs were passaged using Accutase (Sigma-Aldrich A6964) and cryopreserved using BAMBANKER serum-free cell freezing medium (Bulldog Bio BB05). To promote cell survival, ROCK inhibitor (Tocris 1254) was included for single-cell passaging steps but excluded for subsequent media changes.

### Neuronal differentiation

Neuronal differentiation was performed in two steps. In the pre-differentiation step, iPSCs were dissociated and plated on Matrigel (in a T75 or a 10cm^2^ dish) at a density of ∼70,500 cells/cm^2^ in pre-differentiation media (1X Knockout DMEM/F12 [Thermo Fisher Scientific 12660012], 1X NEAA [Thermo Fisher Scientific 11140050], 1X N2 [Thermo Fisher Scientific 17502048], 10 ng/mL NT-3 [PeproTech 450-03], 10 ng/mL BDNF [PeproTech 450-02], 1 μg/mL laminin [Thermo Fisher Scientific 23017-015] supplemented with ROCK inhibitor (Tocris 1254) and 1μg /mL doxycycline. Half media changes (excluding ROCK inhibitor) were performed for two subsequent days. On the third day, the differentiation step began (day 0 of differentiation). Cells were washed once with 1X DPBS (Thermo Fisher Scientific 14190136), dissociated with Accutase (Sigma-Aldrich A6964), and counted. Cells were re-plated into wells pre-coated with 1:2 dilution of poly-L-ornithine (R&D Systems 3436-100-01), at a density of ∼120,000-180,000/cm^2^. For this step, media was changed to differentiation media (0.5X DMEM/F12 [Thermo Fisher Scientific 11330032], 0.5X Neurobasal Plus [Thermo Fisher Scientific A3582901], 1X NEAA [Thermo Fisher Scientific 11140050], 0.5X GlutaMAX [Thermo Fisher Scientific 35050-061], 0.5X N2 [Thermo Fisher Scientific 17502048], 0.5X B27 Plus [Thermo Fisher Scientific A3582801], 10ng/mL NT-3 [PeproTech 450-03], 10ng/mL BDNF [PeproTech 450-02], and 1 μg/mL laminin [Thermo Fisher Scientific 23017-015]). 1μg /mL doxycycline was included in the media on day 0 only. A full media exchange was performed on day 3, followed by a half media exchange on day 7. Cultures were maintained with two media exchanges each week (one full-volume change and one half-volume change).

### Lentiviral infection and library preparation

To produce lentivirus, two 15cm^2^ dishes were seeded with HEK293T cells. The MPRA library was co-transfected with the viral packaging plasmids pMD2.G (Addgene #12259) and psPAX2 (Addgene #12260), using TransIT-293 Transfection Reagent (Mirus Bio 2704). After 10 hours, media was replaced with DMEM containing 5% heat-inactivated FBS supplemented with 1x ViralBoost Reagent (Alstem Bio VB100). After 48 hours, the supernatant was collected and filtered through a 0.5micron PES filter, then concentrated overnight at 4°C with Lenti-X Concentrator (Takara 631231). Virus was then centrifuged at 1,500xg for 45 minutes at 4°C, then the pellet was resuspended in cold 1X DPBS (Thermo Fisher Scientific 14190136) and stored at 4°C.

Virus titration was performed in a 24-well, with 350,000 iNeurons plated per well. Virus was aliquoted into 0μL, 1μL, 2μL, 4μL, 8μL, 16μL, 32μL, and 64μL, with OptiMEM (Thermo Fisher Scientific 31985062) added to bring the total volume to 100μL. To improve iNeuron transduction, 6μl of Viromag R/L (Oz Biosciences RL41000) was added to each virus aliquot, then incubated for 15 minutes at room temperature. Each aliquot was then added dropwise to one well of the 24-well plate. Cells were then placed on a magnetic plate (OZ Biosciences) in the 37°C incubator for 20 minutes, then the magnetic plate was removed. After 7 days, cells were washed once with DPBS and stored at −80°C. Genomic DNA was extracted using a DNeasy kit (Qiagen 69504), then qPCR was performed with 2X PowerUp SYBR Green Master Mix (Thermo Fisher Scientific A25778), with primers to measure the amount of viral DNA (WPRE.F and WPRE.R) relative to genomic DNA (LP34.F and LP34.R)^23^.

For the MPRA, four replicates of day 7 iNeurons (with each replicate comprising 18 million neurons plated across two T75 flasks) were transduced with virus at a MOI ∼30. After one week, flasks were washed with 1X DPBS and frozen at −80°C. RNA and DNA were isolated from each replicate using the Qiagen AllPrep kit (Qiagen 80204). To prepare sequencing libraries, RNA was first treated with TURBO DNase (Thermo Fisher Scientific AM2239) to remove any remaining DNA, and then polyA-tailed mRNA was isolated with oligo d(T) beads (NEB S1550S). 0.7μg of RNA was then reverse-transcribed using SuperScript III RT (Thermo Fisher Scientific 18080093), using the P7-pLSmP-ass16UMI-gfp primer^23^ to add unique molecular identifiers to the cDNA. Finally, library preparation was performed for the cDNA and DNA fractions across two rounds of PCR. The first round added the P5/P7 sequences and unique molecular identifiers, using the entire cDNA fraction or 10μg of DNA as template. Primers were P7-pLSmP-ass16UMI-gfp and P5-pLSmP-5bc-i#^23^. The cycling program was: 98°C for 1 minute, then 3 cycles of 98°C for 10 seconds, 60°C for 30 seconds, and 72°C for 1 minute, with a final incubation of 72°C for 5 minutes. Cleanup was performed with 1.8X HighPrep PCR magnetic beads (Magbio Genomics AC-60050). A second PCR with the same cycling program was performed with P5 and P7 primers, with the cycle number determined through a 10μL qPCR reaction to determine how many cycles were required for the fluorescence to begin to plateau. Final PCR products were purified using 1.8x HighPrep beads, then DNA or RNA replicates were pooled. Finally, DNA and RNA products underwent additional cleanup with gel extraction (NEB T1120S), bead cleanup with 1.8x HighPrep beads, and then were pooled in a 1:3 DNA:RNA ratio. The library was sequenced with an Illumina Novseq 2×250 bp, with 20% PhiX spike in and custom primers (Read 1: pLSmP-ass-seq-ind1, Read 2: pLSmP-bc-seq, index read 1: pLSmP-UMI-seq).

### LentiMPRA analysis

Data analysis was performed using the MPRAflow pipeline^23^ with minor modifications: 1) in the Association workflow, the cigar value of 270M was used as the primary selection criteria to map reads to the reference, and 2) in the Count workflow, merged barcode reads were trimmed to be 15 bp. Only CREs with at least 10 unique barcodes were considered in downstream analysis. The log_2_(RNA/DNA) value for each CRE across each replicate was used for all data analysis. Briefly, a one-way ANOVA with Dunnett’s post hoc test was performed to compare the log_2_(RNA/DNA values) for each of the engineered TF shuffle mutations to its matched reference sequence. A separate ANOVA with Dunnett’s post hoc test was performed for each of the human variants. Each mutated nucleotide was labeled with its numerical position within the entire CRE element to enable binning and comparison with other shuffled sequences. TF hotspots were classified by defining bins with at least 50 shuffled nucleotides to ensure adequate coverage, with 10% of the shuffles significantly decreasing or increasing activity. The full data are available in Supplementary Table 2 and Supplementary Table 6.

### Motif analysis

To discover which motifs contribute most to the activity of each tile, we first extracted all 8–14 nucleotide k-mers from each sequence. To ensure robust statistical testing, we calculated the presence or absence of each k-mer, retaining only those found or missing in at least 20% of the sequences. The association between k-mer presence and expression levels was tested using the non-parametric Brunner-Munzel test, applying a stringent significance threshold of p < 1×10^−3^. Significant k-mers were then matched to known TF motifs using the TomTom^25^ tool with relaxed thresholds (p-value < 0.01), and using both the HOCOMOCO v12 CORE collection and JASPAR as database references. Finally, the expression of the corresponding TFs was then cross-referenced in two iNeuron RNA-seq datasets^26,27^ and the Allen Brain Atlas Brainspan dataset^28^ to prioritize TFs relevant to the cell type and biological system.

### Luciferase assay validation

Gateway cloning technology was used to systematically generate firefly luciferase constructs to validate MPRA results. Briefly, the vector pLS-mP-Luc (Addgene 106253) was digested with XbaI (NEB R0145) and SbfI (NEB R3642). The Gateway reading frame cassette C.1 (ThermoFisher 11828029) was amplified using primers containing overhang pLS-mP-Luc overhang sequence (gateway-cassette-fwd and gateway-cassette-rev). The Gateway adapter was assembled into pLS-mP-Luc by Gibson cloning using the NEBuilder HiFi DNA Assembly kit (NEB E2621). The Gateway compatible pLS-mP-Luc vector was transformed into One Shot *ccdB* Survival *Escherichia coli* cells (Thermo Fisher Scientific 11828029) and grown on chloramphenicol-coated agar plates. Single colonies were grown in liquid culture before plasmid extraction with the NucleoSpin Plasmid Mini kit (Takara 740588.250). Integration of the Gateway cassette was confirmed by Sanger sequencing and digestion with BsrGI (NEB R3575).

The CRE sequence of interest was amplified with BP-containing primers. Then, a BP reaction with Gateway BP Clonase II (Invitrogen 11789020) was performed to recombine the sequence into pDONR, followed by Sanger sequence validation. Point mutations were introduced into the pDONR reference sequence using Q5 Site-Directed Mutagenesis (NEB E0554S). Plasmids were then transformed into *E. coli* with 42°C heat shock and grow overnight on kanamycin-containing LB plates. Colonies were picked into kanamycin-containing LB broth and grown overnight shaking at 37°C. Plasmids were then purified using the NucleoSpin Plasmid miniprep kit (Takara 740588.250) and submitted for Sanger sequencing to verify the sequence. The sequence was then recombined into the firefly destination plasmid using a Gateway LR Clonase II reaction (Invitrogen 11791020), then sub-cloned in *E. coli* as described above. A digest with BsrGI (NEB R3575S) followed by gel electrophoresis was performed to verify proper recombination.

To generate luciferase lentivirus, approximately 250,000 HEK293T cells were seeded in a 24 well plate. After 24 hours, transfection was performed using 0.063 pmol firefly construct, 0.063 pmol of renilla control (Addgene #106292), 108ng pMD2.G (Addgene #12259) and 325 ng psPAX2 (Addgene #12260), with TransIT-293 Transfection Reagent (Mirus Bio 2704) and (OptiMEM (Thermo Fisher Scientific 31985062). After 48 hours, 400μL supernatant was collected and centrifuged at room temperature for 10 minutes at 500xg. 300μL supernatant was then combined with 100μL Lenti-X Concentrator (Takara 631231) and chilled at 4°C overnight. Finally, samples were centrifuged at 4°C for 45 minutes at 1,500xg. Supernatant was discarded and the virus-containing pellet was resuspended in 75μL DPBS.

iNeurons were plated in an opaque white-walled 96-well plate (Thermo Fisher Scientific 165306) at a density of 50,000 cells per well on Day 0 of differentiation. On Day 7, 50μL media was removed from each well and replaced with 50μL of fresh media with 0.3-0.5μL lentivirus. On Day 14, firefly and renilla luciferase signal were read on a Promega GloMax Discover, using the Dual-Luciferase Reporter Assay System (Promega E1960) according to the manufacturer’s instructions.

### iPSC genome editing and clonal expansion

To perform genome editing on WTC11-Ngn2 iPSCs, 30 pmol SpCas9 (Synthego #R20SPCAS9-Sm) was complexed with 150pmol sgRNA (Synthego) at room temperature for 15-30 minutes. Then, 2μg ssODN repair template (IDT) was added, along with 0.5μL GFP plasmid (Lonza). This was nucleofected into 200,000 iPSCs using the P3 Primary Cell Nucleofector X Kit (Lonza V4XP-3032), according to the manufacturer’s instructions and using the pulse setting DN-100. After 24 and 48 hours, GFP signal was evaluated to estimate nucleofection efficiency. Cells were then passaged and genotyped with PCR amplification and Sanger sequencing to estimate cutting and repair efficiency.

To isolate clonal populations, iPSCs were subjected to single-cell sorting with a NanoCellect WOLF Cell Sorter and N1 Single-Cell Dispenser. Before sorting, cells were preconditioned at 37°C for 2 hours with Revitacell supplement (Thermo Fisher Scientific A2644501). Cells were then passaged into single-cell suspension with Accutase (Sigma-Aldrich A6964), diluted to 500,000 cells/mL, and passed through a cell strainer (Corning 352235) before sorting. Single cells were delivered to Matrigel-coated wells of 96-well plate with mTESR plus supplemented with CloneR (StemCell Technologies 05888). Two days after sorting, a full media change was performed with CloneR-supplemented mTESR plus, with another 25% volume CloneR-supplemented media added the next day. After that, media changes were performed every other day. Single-cell clones were expanded, cryopreserved, and genotyped to identify edited clones. DNA was extracted using the DNeasy kit (Qiagen 69504), and the *MECP2* promoter was amplified by PCR using the KAPA HiFi DNA Polymerase Hot Start formulation (Roche 07958889001) with the 5X KAPA HiFi GC Buffer and primers 837_genotyping_fwd and 837_genotyping_rev. Amplicons were screened first by Sanger sequencing to verify knock-in allele editing, then by long-read sequencing to verify the absence of off-target mutations *in cis* along the *MECP2* promoter. Two edited clones and two unedited controls were used for our studies. Clones were tested for mycoplasma contamination with PCR every ten passages. Genome stability was determined by whole genome sequencing with CNV analysis, provided by Medical Genetics Multiomics Laboratory. All four clones (both wild-type and mutant) were negative for any rare (population frequency <1%) CNVs, except for two present in the parental WTC11 genome (a heterozygous deletion in *AUTS2* and a heterozygous duplication in *EFCAB5*).

### shRNA knockdowns

The SGEP vector (Addgene #111170) was digested with XhoI (NEB R0146S) and EcoRI (NEB R3101) and purified using gel extraction with the NEB Monarch DNA Gel Extraction Kit (T1020S). shRNAs were designed using the SplashRNA algorithm^45^. An oligo containing the shRNA and SGEP overhangs was amplified (primers miRE Gib Fwd and miRE Gib Rev) and recombined into linearized SGEP using NEBuilder HiFi DNA Assembly Master Mix (NEB E2621L) at 50°C for 60 minutes. The reaction was then transformed into chemically competent *E. coli* and grown at 37°C overnight on ampicillin-containing LB plates. Colonies were picked, then grown in liquid culture shaking at 37°C overnight. Plasmid DNA was extracted using the NucleoSpin Plasmid Mini kit (Takara 740588.250), then verified by Sanger sequencing. To produce lentivirus, HEK293T cells were seeded in 12-well plates at a density of 800,000 cells per well. The next day, 800ng SGEP was co-transfected with 200ng pMD2.G (Addgene #12259) and 600ng psPAX2 (Addgene #12260), using TransIT-293 Transfection Reagent (Mirus Bio 2704). After 48 hours, supernatant was collected and centrifuged at 500xg for 10 minutes to pellet cell debris. Then, supernatant was concentrated overnight at 4°C with Lenti-X Concentrator (Takara 631231). Virus was then centrifuged at 1,500xg for 45 minutes at 4°C, then the pellet was resuspended in 160μL 1X DPBS (Thermo Fisher Scientific 14190136). iNeurons were transduced at day 7 of differentiation, with 10μL of lentivirus added per 350,000 cells in one well of a 24-well-plate. For *TCF3*, 2.5μL of four independent shRNA lentiviruses were pooled to improve knockdown efficiency.

### RNA and protein analysis

To detect changes in *MECP2* RNA and/or protein expression, 24-well iNeuron plates were washed once with DPBS and then frozen at −80°C on ∼day 28. RNA was harvested using the RNeasy Micro Kit (Qiagen 74004). RNA was reverse transcribed into cDNA using PrimeScript RT Master Mix (Takara Bio RR036A), then diluted to 5ng/μL in water. RT-qPCR was performed using 15ng cDNA, 250nM forward and reverse primers, and 2X PowerUp SYBR Green Master Mix (Thermo Fisher Scientific A25778), with cycling conditions according to the manufacturer’s instructions. Relative gene expression was calculated by ΔΔCt. Statistical significance was calculated using a Welch’s t-test. Primer sequences are in Supplementary Table 7.

To analyze protein, lysis buffer (10mM HEPES pH 7.9, 3mM MgCl_2_, 5mM KCl, 140mM NaCl, 0.1mM EDTA, 0.5% IGEPAL CA-630, 1X phosphatase inhibitor [GenDepot P3200], 1X protease inhibitor [GenDepot P3100], and 1:2500 nuclease [Thermo Scientific 88702]) was added to iNeuron plates thawed on ice. Cells were lysed on ice with occasional tapping until neuronal network had lifted, then cells and supernatant were transferred to a 1.5mL Eppendorf tube and centrifuged at max speed at 4°C for 15 minutes. The lysate was transferred to a new tube and protein concentration was quantified using a BCA assay (Thermo Fisher Scientific 23225). 10μg of total protein as mixed with 1X NuPAGE LDS Sample Buffer (Thermo Fisher Scientific NP0008) and 5% beta-mercaptoethanol (Bio-Rad 1610710), then boiled at 95°C for 10 minutes. Prepared samples were run on a NuPAGE 4-12% Bis-Tris gradient gel with MOPS SDS running buffer (Boston BioProducts BP-178). Separated proteins were transferred to a 0.2μm nitrocellulose membrane (Cytiva 10600004) using a constant 0.25mA applied at 4°C for 2 hours in a 1X Tris-Glycine transfer buffer with 10% methanol. Membranes were then blocked in 1X Intercept (TBS) Blocking buffer (LICORbio 927-60025) for 1 hour at room temperature. Membranes were then probed with primary antibodies in 0.5X Blocking Buffer (diluted with 1X TBST) at 4°C overnight. The following antibodies and dilutions were used: anti-MeCP2 (rabbit monoclonal D4F3, Cell Signaling Technologies 3456), anti-Vinculin (mouse monoclonal hVIN-1, Sigma-Aldrich V9131). Membranes were washed three times with TBST for 5 minutes, then incubated in LiCOR secondary antibodies (1:20,000 dilution) in 0.5x Blocking Buffer for one hour. Membranes were washed three times with TBST for 5 minutes, then imaged on a LiCOR Odyssey M imager. Band intensity was quantified using Licor Image Studio software. Statistical significance was calculated using a Welch’s t-test.

### CUT&Tag

iNeurons were collected on Day 24 of differentiation for CUT&Tag library preparation. The EpiCypher CUTANA Direct-to-PCR CUT&Tag Protocol was followed with few modifications. iNeurons were washed once with PBS, then lysed in Nuclei Extraction Buffer on ice. Cells were collected and triturated to reduce clumping, before centrifugation and continuation of the protocol. Two wild-type clones and two chrX:154097837 A>G clones were used, with three technical replicates per reaction. Each reaction used 100,000 nuclei and 0.5μg of antibody (NFYB [Diagenode C15410241] or IgG [EpiCypher 13-0042]). To perform the index PCR, NFY-B and IgG samples were amplified for 18 cycles. Libraries were pooled and sequenced to a depth of 20M reads per library on a NovaSeq X Plus.

Adaptors were trimmed from fastq reads using cutadapt (-a CTGTCTCTTATACACATCT -A CTGTCTCTTATACACATCT --nextseq-trim=30, --minimum-length 20). Trimmed reads were aligned to hg38 using bowtie2 (--local --very-sensitive --no-mixed --no-discordant --phred33). Output files were converted into bigwig files to visualize in the Integrative Genomics Viewer (IGV) using the deepTools bamCoverage function (--centerReads –ignoreDuplicates –normalizeUsing none). Peaks were called from BAM files using SEACR with a top 1% signal cut-off.

To evaluate global NFY-B binding patterns, consensus peaks were identified using bedtools, first merging overlapping peaks and then intersecting across replicates to identify peaks present in at least 8 of the 12 samples. These were compared to NFY-B ChIP-Seq datasets from ENCODE (ENCFF552QUI, ENCFF156MUM, and ENCFF718ZFY), using bedtools intersect to count how many peaks were shared with the other three datasets. To test the iNeuron consensus peaks for promoter enrichment consistent with known NFY binding patterns^33^, Genomic Association Tester (GAT)^46^ was used to compare the observed distribution of iNeuron consensus peaks to a null distribution generated by 1,000 random peak redistributions that conserved interval length and chromosome distribution. Promoter regions were defined using GENCODE promoter windows (1000 bp immediately upstream of the MANE Select TSS), gene bodies were defined as the span of GENCODE primary transcripts, and intergenic regions were defined as all remaining genomic space. P-values were computed as one-sided tail probabilities, followed by Benjamini-Hochberg multiple-testing corrections. For identifying enriched motifs in the iNeuron consensus peak set, we used the motif discovery software HOMER^47^.

To rank NFY-B signal at promoters, bedtools was used to intersect the consensus iNeuron peaks with GENCODE promoter windows. Then, multiBigwigSummary was used to calculate the mean NFY-B signal from bigwig files across these promoter coordinates for each replicate. Promoter-level NFY-B signal values were ranked within each replicate, and percentile ranks were calculated for the *MECP2* promoter within each replicate.

## Data Availability

Whole-genome sequencing data from the Centers for Common Disease Genomics is available on anvilproject.org. The MPRA data, including all relevant sequences, their barcode counts, and their associated log_2_(RNA/DNA) values, are available in Supplementary Tables 2 and 6. The NFY-B CUT&Tag data has been deposited to GEO (Accession GSE330364).

## Author contributions

R.M-S, Y.S., E.E.E., and H.Y.Z conceptualized the study. R.M-S, F.C., and Y.L. performed experiments. Y.S., T.W., and E.E.E. performed variant discovery and analysis. A.P-K. performed motif analysis, and Y.Z. and Z.L. provided computational support. K.H., D.K., Y.S., H. B-R., A.C., T.N., E.E.E., and H.Y.Z. supported experimental planning and data interpretation. R.M-S. and H.Y.Z. wrote the manuscript with contributions and feedback from all authors.

