## Supplementary Figures for "A massively parallel reporter assay of *MECP2* cis-regulatory elements reveals genetic candidates for male-biased autism"

Supplementary Information

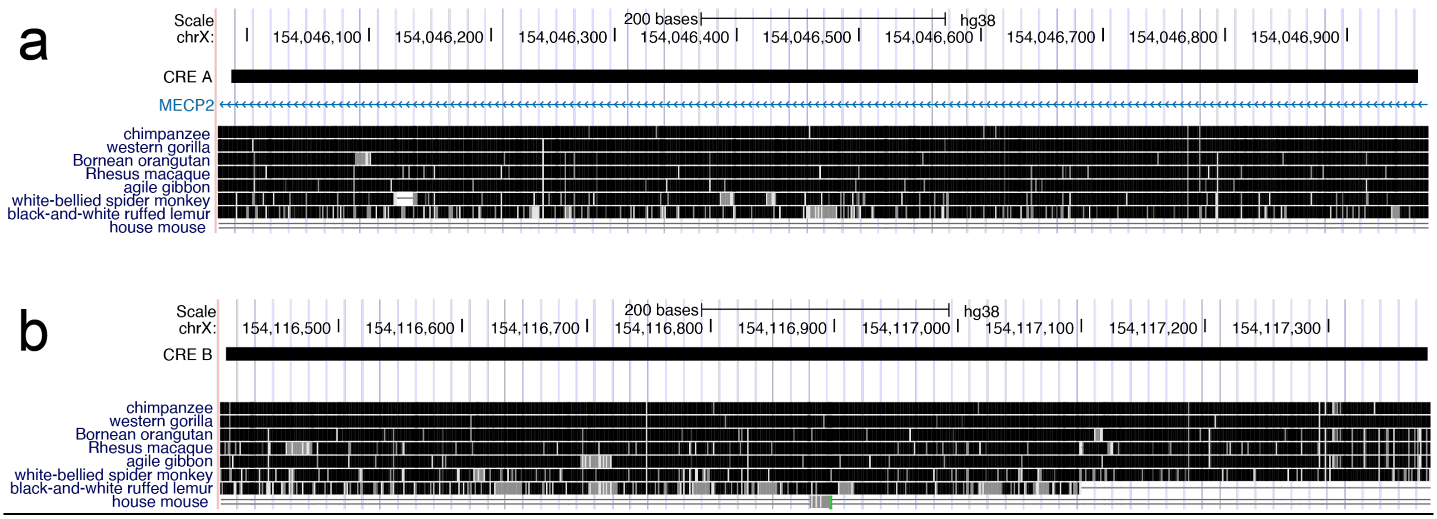


**Figure S1. CREs A and B are conserved across nonhuman primates.** Multiple species alignment for CRE A **(a)** and CRE B **(b)**. Selected tracks from the UCSC Browser Zoonomia+Primates 447 Cactus alignment are shown.

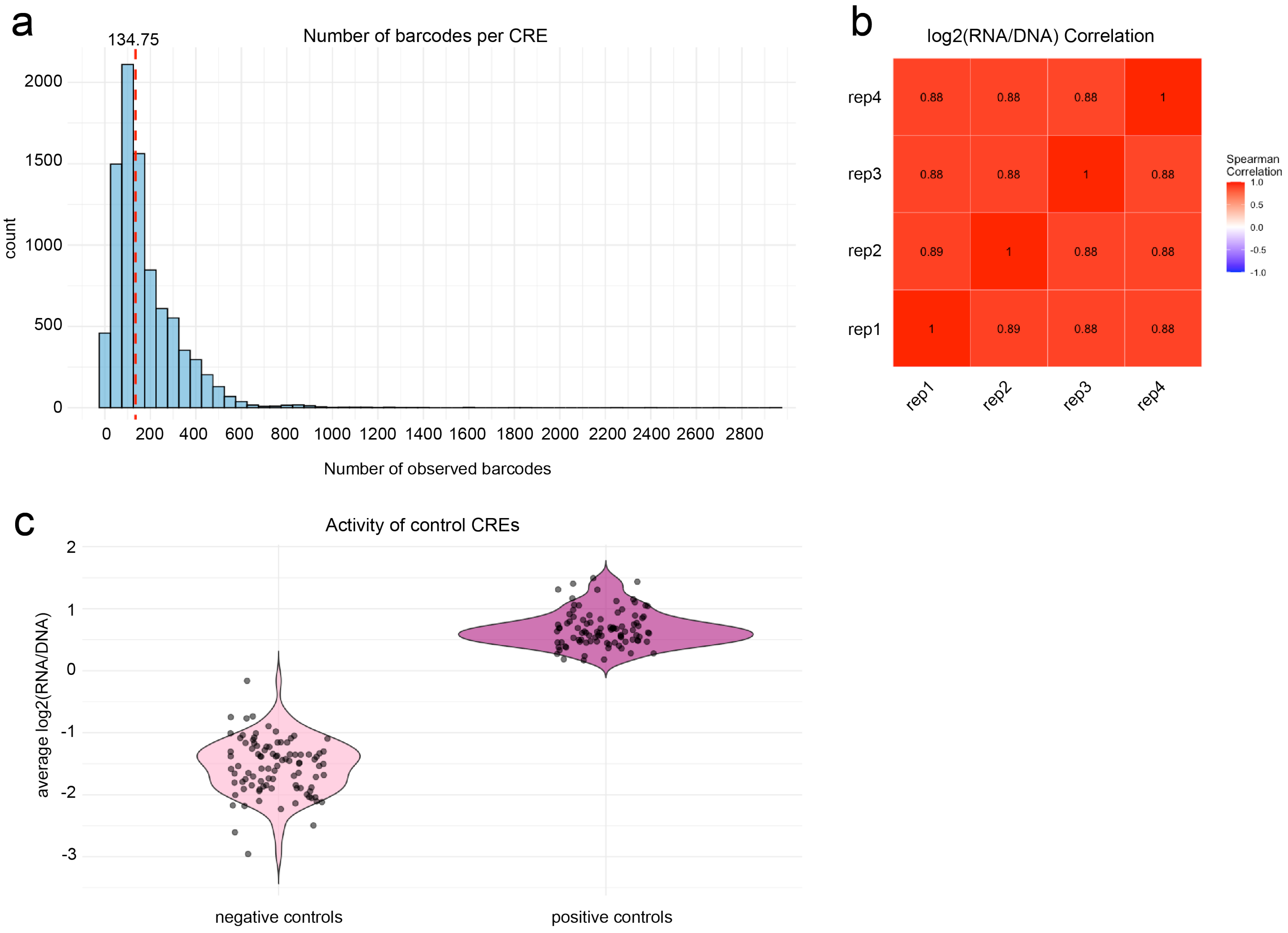


**Figure S2. MPRA quality control metrics. (a)** Distribution of the number of barcodes per CRE, with the majority of CREs associated with 50-300 independent barcodes. **(b)** Spearman correlation of the $\log_2$(RNA/DNA) values for each tested sequence across each of the four tested replicates. **(c)** Mean $\log_2$(RNA/DNA) value for each of the negative and positive controls included in the library.

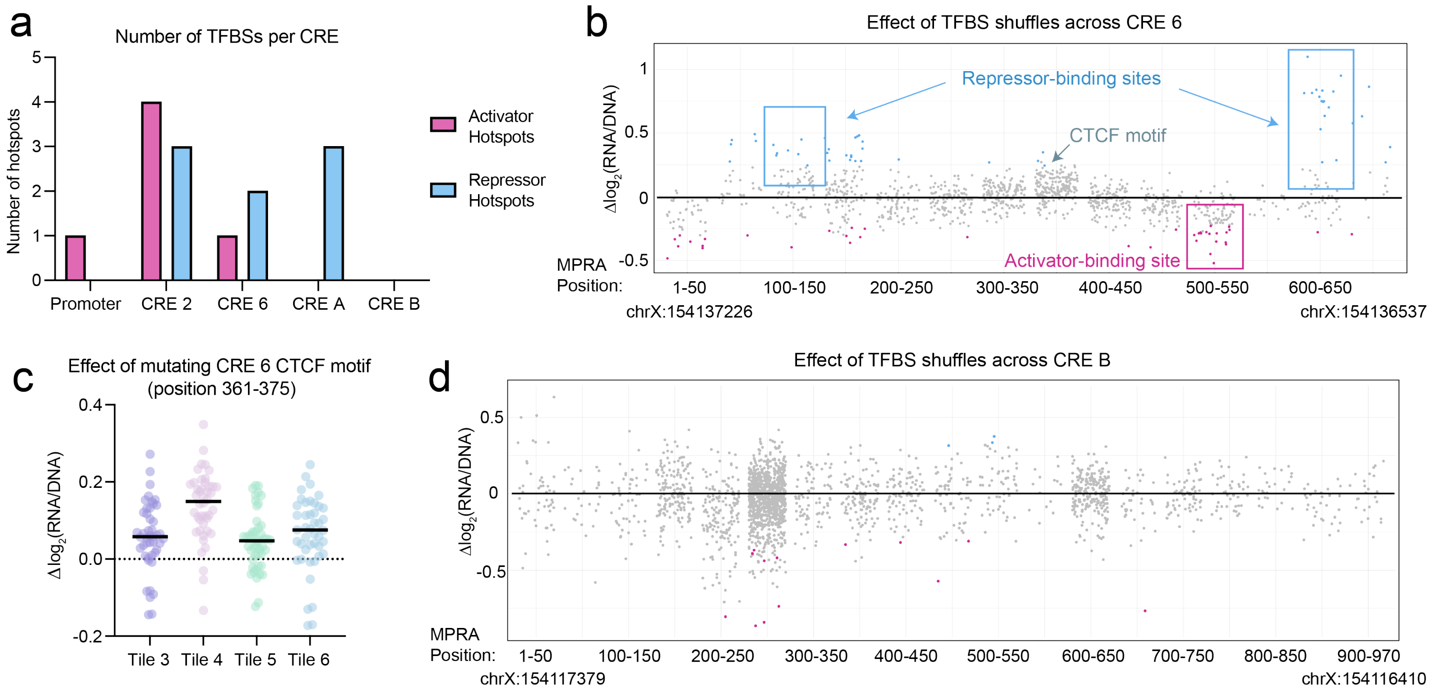


**Figure S3. Defining activator and repressor hotspots. (a)** Number of activator and repressor hotspots identified across all five CREs. **(b)** The effect of shuffling TFBSs in CRE 6. Statistically significant ($p_{adj}$<0.05) values are shown in blue or pink. The pink or blue boxes denote activator- or repressor-binding sites, respectively. **(c)** The effect of shuffling a central CTCF motif on MPRA activity in four different tiles, with the mean value indicated by a black line. **(d)** The effect of shuffling TFBSs in CRE B.

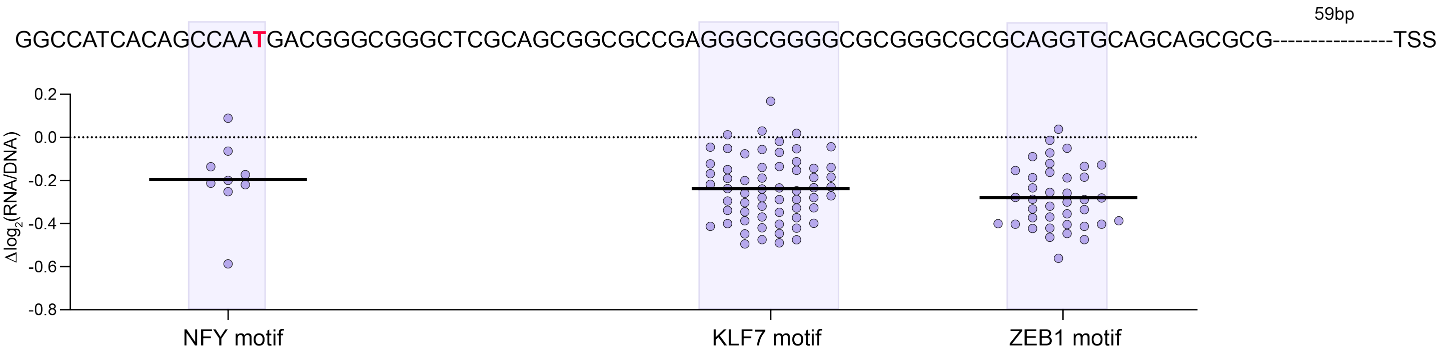


**Figure S4. TFBSs in the *MECP2* proximal promoter.** Critical sequence at the 3' end of the proximal promoter, with three TF motifs highlighted in purple and the effect of shuffling these motifs shown below. Only shuffles that mutate the motif at least two times are included. The -7837 residue within the NFY motif is highlighted in red.

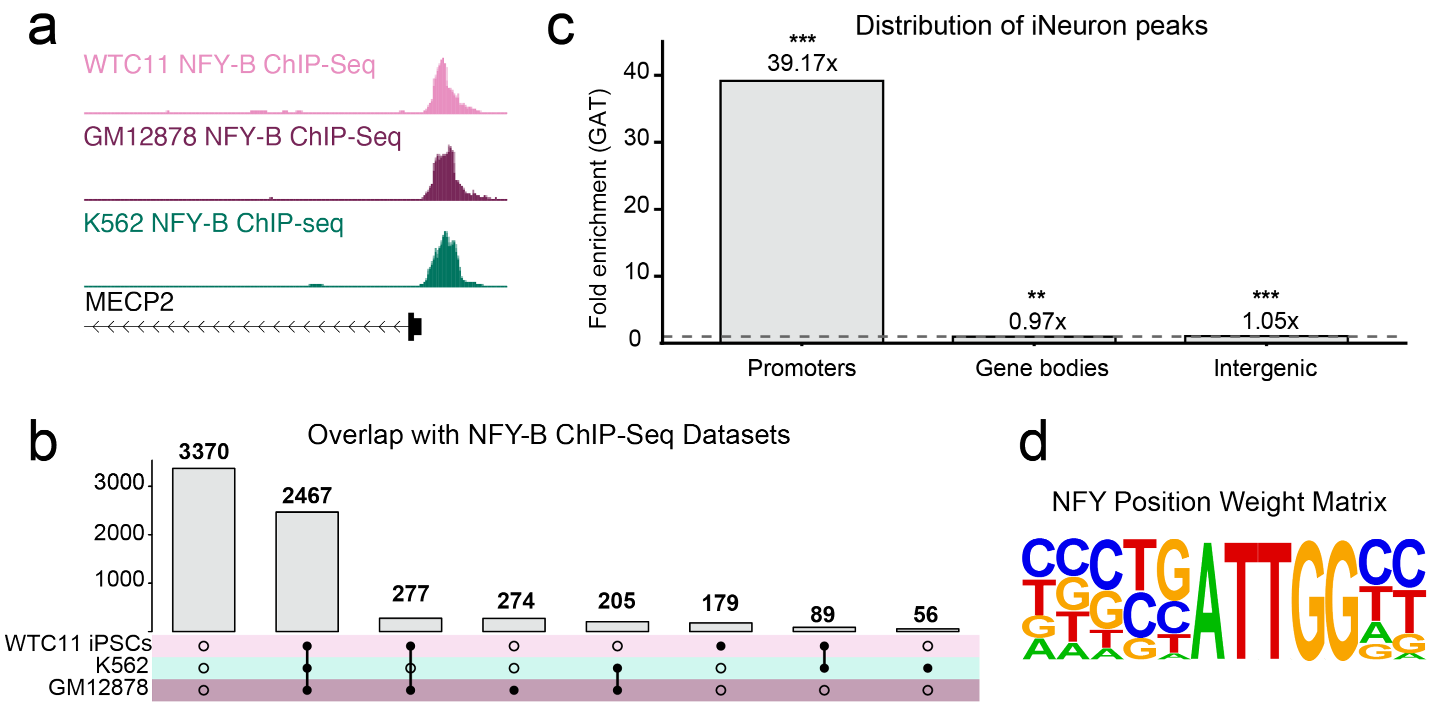


**Figure S5. Characteristics of NFY-B CUT&Tag peaks. (a)** NFY-B signal at the *MECP2* promoter from three ENCODE ChIP-Seq datasets. **(b)** Upset plot showing the number of shared NFY-B peaks between iNeuron CUT&Tag (this study) and three ENCODE ChIP-Seq datasets in diverse cell types. **(c)** GAT enrichment analysis of iNeuron peaks in promoters, gene bodies, and intergenic regions. *** p<0.001, ** p<0.01. **(d)** Results of *de novo* motif discovery for iNeuron peaks ($p=1x10^{-1026}$).
